# Response to Tcherkez and Farquhar: Rubisco adaptation is more limited by phylogenetic constraint than by catalytic trade-off

**DOI:** 10.1101/2023.01.07.523088

**Authors:** Jacques W. Bouvier, Steven Kelly

## Abstract

Rubisco is the primary entry point for carbon into the biosphere. It has been widely proposed that rubisco is highly constrained by catalytic trade-offs due to correlations between the enzyme’s kinetic traits across species. In previous work, we have shown that these correlations, and thus the strength of catalytic trade-offs, have been over-estimated due to the presence of phylogenetic signal in the kinetic trait data (Bouvier et al., 2021). We demonstrated that only canonical trade-offs between the Michaelis constant for CO_2_ and carboxylase turnover, and between the Michaelis constants for CO_2_ and O_2_ were robust to phylogenetic effects. We further demonstrated that phylogenetic constraints have limited rubisco adaptation to a greater extent than the combined action of catalytic trade-offs. Recently, however, our claims have been contested by Tcherkez and Farquhar (2021), who have argued that the phylogenetic signal we detect in rubisco kinetic traits is an artefact of species sampling, the use of *rbcL-based* trees for phylogenetic inference, laboratory-to-laboratory variability in kinetic measurements, and homoplasy of the C_4_ trait. In the present article, we respond to these criticisms on a point-by-point basis and conclusively show that all are either incorrect or invalid. As such, we stand by our original conclusions. Specifically, the magnitude of rubisco catalytic trade-offs have been overestimated in previous analyses due to phylogenetic biases, and rubisco kinetic evolution has in fact been more limited by phylogenetic constraint.

## Introduction

Phylogenetic signal is the phenomenon by which closely related individuals resemble each other more than individuals that are distantly related (Blomberg et al., 2003; Blomberg and Garland, 2002). In biological data, the presence of phylogenetic signal is determined by evaluating the covariation in traits across species in the context of their proximity to one another on a phylogenetic tree. Although such a phylogenetic structure in interspecific trait variation might be expected *a priori* based on the action of inheritance and descent with modification, phylogenetic signal is not universal in traits or across taxonomic groups (Blomberg et al., 2003; Freckleton et al., 2002; Kamilar and Cooper, 2013; Losos, 2008). For instance, many evolutionary processes including rapid adaptation, divergent selection, and convergent evolution can lead to significant departures from phylogenetic signal by either causing similarity among species to be independent of evolutionary distance, or by causing distantly related species to appear more similar on average compared to those which share more recent common ancestry (Hansen and Martins, 1996; Kamilar and Cooper, 2013; Kelly, Grenyer and Scotland, 2014). Moreover, although genetic sequence similarity is used to infer evolutionary history among species, it can be difficult to predict the molecular basis of trait variation. In enzymes, for example, single point mutations can have dramatic consequences for catalytic function (Cleton-Jansen et al., 1991; Johnson et al., 2001; Villar et al., 1997) whilst divergent sequences can maintain similar biochemical properties (Espadaler et al., 2008). As such, studying phylogenetic signal can be instructive to interpret patterns of biological diversity and their underpinning evolutionary processes, as well as to help map the sequence–function landscape to better understand genotype–phenotype interactions.

Aside from the fundamental interest in examining the phylogenetic basis of trait data, quantification of phylogenetic signal is also of importance in comparative biology. This is because the shared ancestry of all taxa as determined by the hierarchical tree of life causes a “serious statistical problem” (Felsenstein, 1985) in cross-species analyses. Specifically, this problem arises due to the fact that biological datapoints are not independent observations, as is a core assumption of conventional statistical methods (table 1). Instead, the phylogenetic non-independence between taxa means that the trait variation captured by a given dataset is an artefact of species sampling, where the magnitude of this introduced sampling bias is affected by the strength and pattern of phylogenetic signal in the traits of interest. However, although this non-independence of species violates the assumptions of conventional statistical methods, specialist phylogenetically aware methods have been developed to circumvent this issue (Freckleton et al., 2002; Lindenfors et al., 2010; Motani and Schmitz, 2011; Revell, 2009). In contrast to their non-phylogenetic counterparts, these phylogenetic methods do not assume independence among data points, but instead measure and account for the relative non-independence of all species in the context of their phylogenetic relatedness (table 1). In this way, the invention of phylogenetic statistical tools has enabled biologically meaningful conclusions to be drawn from cross-species analyses.

**Table 1.** Overview of the key differences between conventional non-phylogenetic generalized least squares regression models and phylogenetic generalised least squares regression models.

|  | Assumes independence? | Data Requirements |
| --- | --- | --- |
| Non-phylogenetic regression | ✓ Assumes datapoints are independent and randomly sampled | ✓ Requires comparative trait data<br>☐ Does not require phylogenetic information (datapoints are assumed to be independent and not related in phylogenetic space) |
| Phylogenetic regression | ☐ Does not assume independence among datapoints | ✓ Requires comparative trait data<br>✓ Requires phylogenetic information to measure and correct for the phylogenetic signal (and phylogenetic non-independence) of species datapoints |

One of the most common phylogenetic statistical methods, the phylogenetic generalized least squares regression (Grafen, 1989; Martins and Hansen, 1997; Pagel, 1999, 1997; Rohlf, 2001), assesses the association between continuous biological traits using an analogous approach to a conventional least squares regression. However, in contrast to a conventional regression which assumes independence and thus weights all datapoints equally, the phylogenetic regression works by weighting the importance of data differentially depending on the degree of shared ancestry of species with one another (Figure 1). For an excellent review of the conceptual and technical background to this tool, see (Symonds and Blomberg, 2014). In brief, this feat is achieved by incorporating a phylogenetic parameter in the model formulation which accounts for the phylogenetic signal in the residual errors of the data (i.e., the co-variation between species relatedness and their position in X~Y trait space). If no phylogenetic signal exists in the residual errors, the phylogenetic parameter of the model is equal to 0 and a standard non-phylogenetic regression is performed (Figure 1). In contrast, as the phylogenetic signal in the residual errors increases, this is accordingly controlled for in the model by reducing the weight of data from groups of closely related species on the phylogenetic tree as compared to species which are more evolutionarily distant (Figure 1). Therefore, in this way, phylogenetic least squares regression accounts for the non-independence of biological data and generates a robust and accurate estimate of the gradient, intercept, coefficient of variation (*R*^2^) and associated p-values for biological associations (Figure 1).

**Figure 1.**
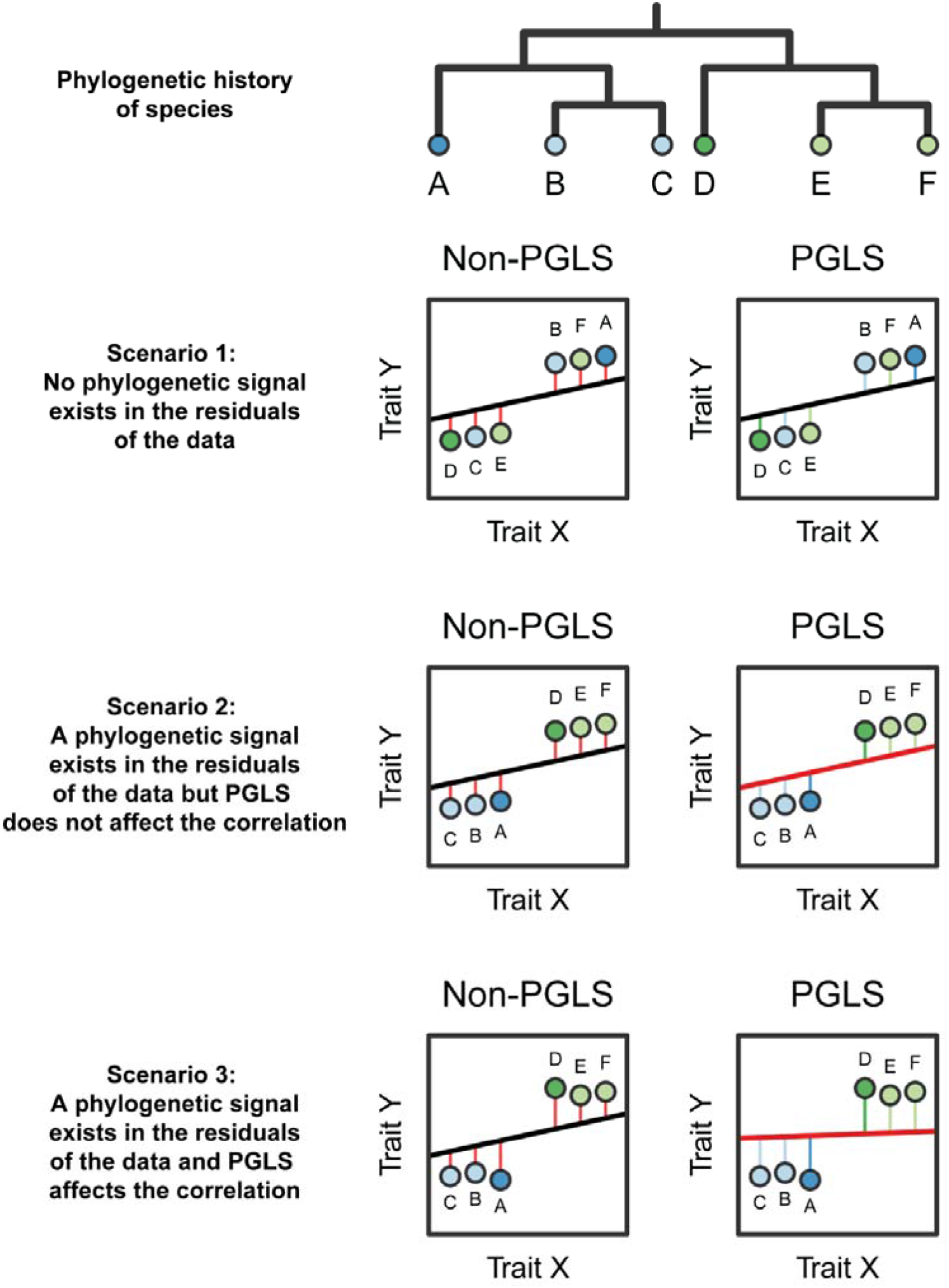
Cartoon to illustrate how conventional and phylogenetic generalized least squares regression models infer relationships between biological traits. Under all scenarios 1 – 3, conventional non-phylogenetic regression models are blind to the underlying phylogenetic history of the sampled species and compute the relationship between trait X and trait Y assuming all datapoints are independent observations. In contrast, phylogenetic regression models measure the co-variation between species relatedness and their position in X~Y trait space, subsequently taking this information into account during downstream model fitting. Specifically, if there is no phylogenetic signal in the residuals of the data, a phylogenetic correction of the data is not necessary and a conventional regression model will be fitted (Scenario 1). Alternately, if a phylogenetic signal in the residual errors is detected, a phylogenetic correction of the data will be performed where depending on the data, this may or may not affect the observed relationship relative to if a non-phylogenetic regression model was fitted (Scenario 2 and 3). These scenarios are illustrated using trait data from six species (identified by letters A—F). The evolutionary relationships among these species are described based on the phylogenetic tree and are also visualized by the colours of each species datapoint. Lines of best fit from conventional and phylogenetic generalized least squares regression models are shown in black and red, respectively.

The necessity to consider phylogenetic context when analyzing interspecies data has been widely adopted in many fields of biology, including ecology (e.g., Wu et al., 2021), animal behavior (e.g., Balasubramaniam et al., 2012), anthropology (e.g., Lukas, Towner and Borgerhoff Mulder, 2021) and conservation science (e.g., Fritz and Purvis, 2010). In contrast, in other sub-disciplines of biology, especially those which study phenomena at the cell or molecular level, phylogenetic statistical practices are less commonplace. Thus, it is likely that many studies which have compared quantitative data in cellular or molecular datasets, particularly those which do so between species, have been affected by the presence of phylogenetic signal.

In a recent study, we performed a cross-species analysis of the kinetic traits of the enzyme rubisco (ribulose-1,5-bisphosphate [RuBP] carboxylase/oxygenase). Specifically, we set out to examine the constraints which have limited the enzyme’s adaptation whilst correctly accounting for the phylogenetic signal arising from the non-independence of the kinetic measurements (Bouvier et al., 2021). Rubisco presents an interesting subject as the basis for such investigation, as despite being the principal carbon-fixing enzyme in the biosphere (Field et al., 1998; Tabita et al., 2008), it is widely considered to be poorly optimised because it exhibits a modest rate of CO_2_ turnover (Badger et al., 1998; Bar-Even et al., 2011) and catalyses a mostly counterproductive reaction with O_2_ (Bowes et al., 1971; Chollet, 1977; Sharkey, 2020). The initial hypothesis which was put forward to explain this paradox of why this enzyme of paramount biological importance appears poorly adapted for its role in primary CO_2_ fixation was pioneered by two comparative studies which reported severe antagonistic correlations between rubisco kinetic traits across species and proposed that these were caused by chemical constraints on the catalytic mechanism of the enzyme (Savir et al., 2010; Tcherkez et al., 2006). However, these studies, as well as all other subsequent analyses which have investigated rubisco kinetic trait correlations (Flamholz et al., 2019; Iñiguez et al., 2020; Young et al., 2016), failed to consider the phylogenetic context of the species being analysed and the fact that all existing rubisco are related by evolution from a common ancestor. As such, it is possible that these correlations suffered from Felsenstein’s “serious statistical problem” (Felsenstein, 1985) meaning that they are potentially an artefact of the presence of phylogenetic signal in rubisco kinetics and the non-independence of species on the phylogenetic tree.

In Bouvier et al., (2021), we found that all of rubisco’s kinetic traits exhibit strong phylogenetic signal (table 2). This signal was observed when analysing the kinetic data across the tree of life, including among C_3_ angiosperms, among all angiosperms, and among all photosynthetic organisms for which kinetic data were available, respectively (table 2). Given this observed non-independence of the kinetic data, we re-evaluated the kinetic trait correlations between species using phylogenetic least squares regression and found that all were attenuated compared to previously published values (Figure 2A – 2C) (Bouvier et al., 2021). Thus, phylogenetic non-independence had caused catalytic trade-offs to be over-estimated in previous analyses. Despite this, we showed that there was nevertheless still moderate antagonism between the Michaelis constant for CO_2_ (*K*_C_) and carboxylase turnover (*k*_catC_) (variance explained = 21–37%), and between the Michaelis constants for CO_2_ and O_2_ (*K*_O_) (variance explained 9–19%), though, all other catalytic trade-offs were negligible or non-significant (Figure 2B and 2C) (Bouvier et al., 2021). Following this, we demonstrated that phylogenetic constraints explained more variation in rubisco kinetics (variance explained 30–61%), and have thus had a larger impact on limiting enzyme adaptation, compared to the combined action of all catalytic trade-offs (variance explained 6–9%) (Bouvier et al., 2021). In summary, therefore, although rubisco catalytic trade-offs exist, the strength of these trade-offs are weaker than previously thought and represent a minor component in limiting rubisco adaptation compared to phylogenetic constraints.

**Figure 2.**
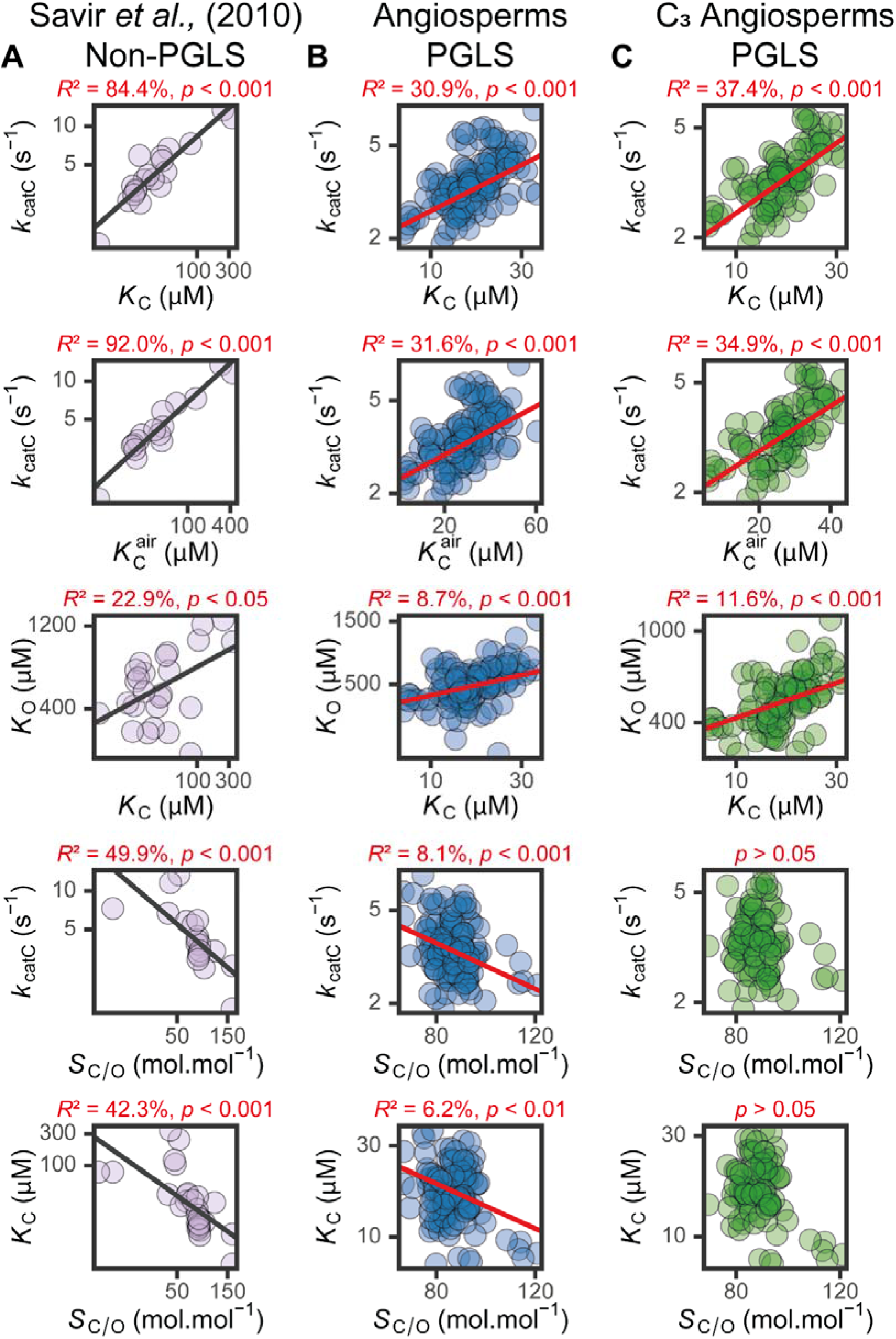
Canonical kinetic trait trade-offs in rubisco as estimated using conventional and phylogenetic generalized least squares regression models. **A)** The pairwise correlations between rubisco kinetic traits across diverse photosynthetic organisms as proposed in the seminal cross-species analysis of (Savir et al., 2010), measured using non-phylogenetic generalized least squares regression and a small dataset of species. **B)** The pairwise correlations between rubisco kinetic traits in angiosperms as described in (Bouvier et al., 2021), measured using phylogenetic generalized least squares regression. **C)** As in (B) but including only C_3_ angiosperms. Correlation coefficients (R^2^) and associated P-values between rubisco kinetic trait correlations are displayed above each respective plot. Significance values are represented as α levels, where; α□=□0.001 if P□<□0.001, α□=□0.01 if 0.001□<□P□<□0.01, α□=□0.05 if 0.01□<□P□<□0.05, and α = ns if P□>□□.05. Both x and y axes for all plots are on a log transformed scale and respective units are shown in axis labels. Lines of best fit from conventional and phylogenetic generalized least squares regression models are shown in black and red, respectively.

**Table 2.**
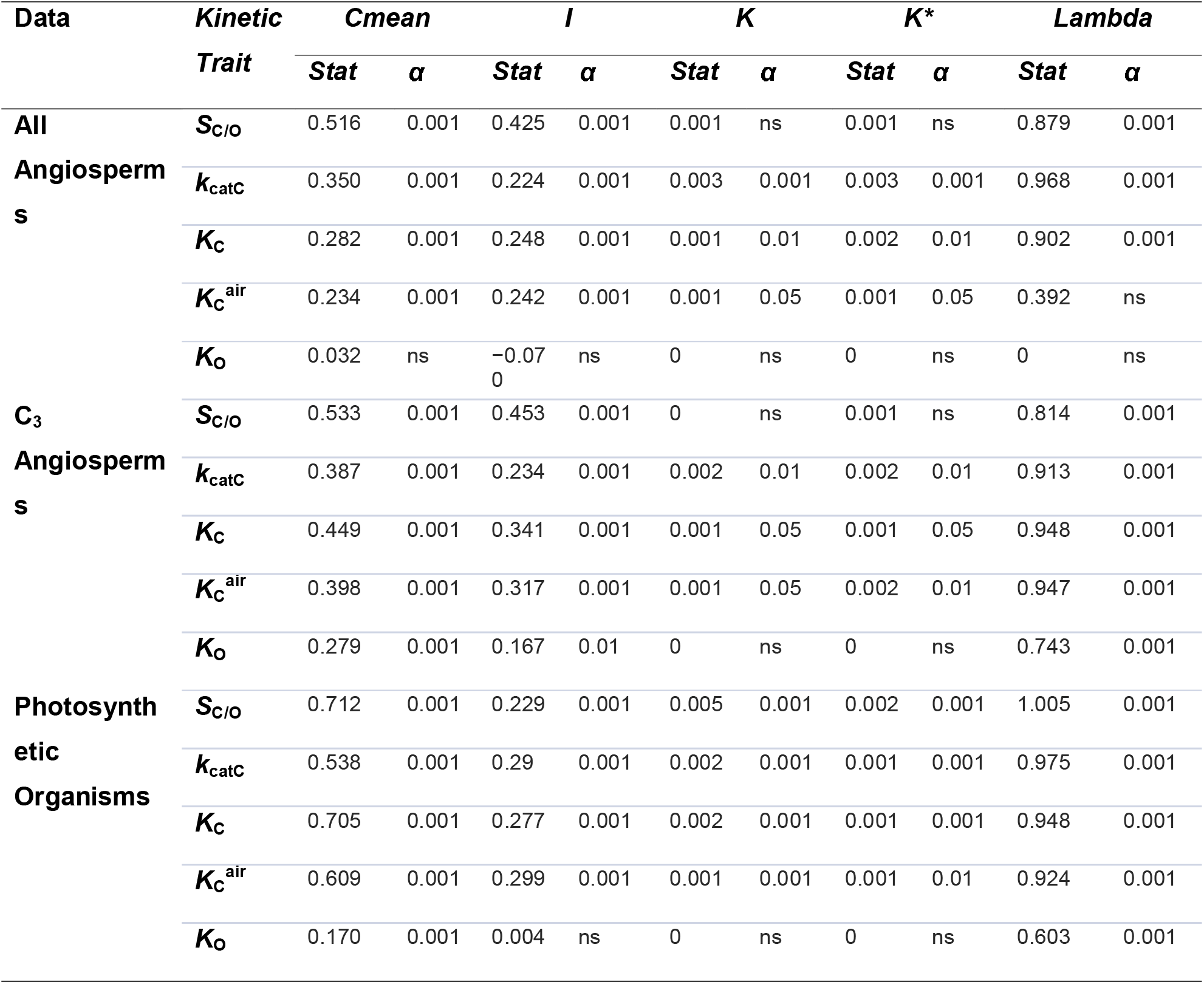
The phylogenetic signal strength and associated significance level in rubisco kinetic traits using five different signal detection methods, as reported in (Bouvier et al., 2021). Data are shown from the analysis of angiosperms, C_3_ angiosperms, and all studied photosynthetic organisms, respectively. NOTE. Statistics are rounded to three decimal places and significance values are represented as α levels, where α□=□0.001 if P□<□0.001, α□=□0.01 if 0.001□<□P□<□0.01, α□=□0.05 if 0.01 □<□P□<□0.05, and α = ns if P□>□0.05.

Although the above work sheds light on our understanding of the constraints that have shaped rubisco adaptation, the validity of the results presented in Bouvier et al., (2021) have been brought into question in a recent opinion piece (Tcherkez and Farquhar, 2021). In the present study, we address and refute the criticisms made by Tcherkez and Farquhar. In doing so, we confirm that the data we previously presented is valid and robust and thus our original conclusions are unaltered. Specifically, although catalytic trade-offs are present in rubisco as determined by chemistry, the extent of these constraints on enzyme adaptation is overestimated unless phylogenetically appropriate methods are used. Instead, phylogenetic constraints have provided a more severe limitation on rubisco optimisation.

## Results

### Kinetic and phylogenetic data

As the basis of the analysis in the present study, the same rubisco kinetic dataset was used as described in Bouvier et al., (2021). This includes experimental measurements of CO_2_/O_2_ specificity (*S*_C/O_). maximum carboxylase turnover rate per active site (*k*_catC_), the respective Michaelis constant (i.e., the substrate concentration at half-saturated catalysed rate) for both CO_2_ (*K*_C_) and O_2_ (*K*_O_) substrates, as well as an inferred measurement of the Michaelis constant for CO_2_ in 20.95% O_2_ air (*K*_C_^air^) calculated from the formula *K*_C_^air^ = *K*_C_ + (*K*_C_ [O_2_]/*K*_O_), where 20.95% [O_2_] in water is 253 μM. As before, statistical interrogation of this dataset was restricted to log transformed trait values to ensure all data conformed to the distribution assumptions of the statistical analysis herein. Moreover, consistent with the analysis in Bouvier et al., (2021), the same phylogenetic trees of studied species were used (unless explicitly stated) as previously generated from the coding sequences of the rubisco large subunit (*rbcL*) gene.

To assess whether rubisco in different angiosperms display similar kinetics as a consequence of their phylogenetic relationship, the presence and magnitude of phylogenetic signal in rubisco kinetic traits was assessed using the statistical tools previously applied in Bouvier et al., (2021), including Pagel’s lambda (Pagel, 1999), Blomberg’s *K* (Blomberg et al., 2003), Blomberg’s *K*\* (Blomberg et al., 2003), Moran’s *I* (Gittleman and Kot, 1990) and Abouheif’s *Cmean* (Abouheif, 1999). For an overview of the inherent differences between these methods utilized see (Bouvier et al., 2021), or for more extensive discussion see (Münkemüller et al., 2012). However, in brief, signal strength for each metric typically varies between 0 (absence of phylogenetic signal) and 1 (strong phylogenetic signal), and we assess the presence or absence of a phylogenetic signal in each trait as the majority result across these methods (i.e., the consensus result in ≥3 out of 5 methods tested).

### A phylogenetic signal cannot be generated with a randomly simulated arbitrary trait

The first criticism of our work made by Tcherkez and Farquhar (2021) states that a phylogenetic signal may be generated using an arbitrary trait that is randomly distributed across species. However, the presence of phylogenetic signal in Bouvier et al., (2021) was assessed using a combination of robust statistical methods based on both non-parametric permutation tests (Blomberg’s *K*, Blomberg’s *K*\*, Moran’s *I* and Abouheif’s *Cmean*) as well as likelihood ratio tests (in the utilized implementation of Pagel’s lambda). In brief, permutation tests evaluate the probability that the null hypothesis of absence of signal is true by comparing the estimated test statistic for a given dataset to the distribution of values of this statistic obtained when randomly re-shuffling the data among species on the tree across a set number of replicates (*n* = 10,000 in Bouvier et al., (2021)). Conversely, likelihood ratio tests evaluate the probability that the null hypothesis is true by comparing the ratio of the log likelihood of the null hypothesis to that of the alternate hypothesis using the formula −2[log likelihood (null hypothesis) – log likelihood (alternative hypothesis)]. In both cases, if the computed p-value satisfies the given threshold of significance, the null hypothesis is rejected, and phylogenetic signal is deemed to be statistically significant. Specifically, in Bouvier et al., (2021), the majority of p-values reported for phylogenetic signal in rubisco kinetic traits were below p = 0.001 across the suite of detection methods used (table 2). This means that in these cases, there is less than a 0.1% chance of observing a similar or stronger phylogenetic signal by chance. Thus, to put in plainly, it is not true that an arbitrary trait can spuriously generate a phylogenetic signal in a reliable manner. In fact, by definition, an arbitrary trait would produce a similar signal in fewer than 1 in 1,000 simulations. Nevertheless, although these statistical methods are powerful and perform well under a range of scenarios, we reaffirm below using two independent analyses that an arbitrary trait cannot reliably produce a phylogenetic signal. Thus, the above statement on this matter made by Tcherkez and Farquhar (2021) is false.

First, to confirm the validity of our implemented phylogenetic tests and demonstrate that arbitrary traits cannot produce a statistically significant phylogenetic signal, we subject 100,000 replicates of a randomized assignment of kinetic trait data across species on the rubisco tree to phylogenetic signal analysis (i.e., recapitulating the methods of the permutation-based signal tests manually). Here, when using a cut-off threshold of significance at p ≤ 0.05, interrogation of these randomized data revealed a significant signal could only be detected in 5% of the re-shuffled kinetic trait data when considering the majority of signal detection methods (i.e., exactly matching the proportion that would be predicted to occur by chance), except Pagel’s lambda which exhibited a considerably lower significance rate (Supplemental File 1, table S1). This result was similarly observed in the analysis across all angiosperms, as well as across the subset of C_3_ angiosperms (Supplemental File 1, table S1). In total, this resulted in a type I (false positive) error rate in the randomized data at between 1.1 and 1.7% when taking the consensus result across at least 3 out of 5 phylogenetic signal detection methods (Supplemental File 1, table S1).

To further demonstrate that any arbitrary trait cannot generate a phylogenetic signal, 100,000 datasets were also randomly simulated *de novo* and subject to phylogenetic signal interrogation as above. To achieve this, we used the rubisco phylogenetic tree in Bouvier et al., (2021) and replaced the kinetic data for each species with randomly generated data. For this purpose, the mean and standard deviation of the real kinetic traits were used to guide the simulated data such that 100,000 arbitrary variables were produced with the same distributional properties as each trait. Analogous to above, when using a cut-off threshold of significance at p ≤ 0.05, we observe a phylogenetic signal in 5% of all arbitrary traits in both the analysis across all angiosperms, as well as across the subset of C_3_ angiosperms for the majority of detection methods (i.e., again corresponding to that which would be expected by chance) (Supplemental File 1, table S2). Accordingly, as above, the type I false positive rate was observed at between 1.4 and 1.6% when considering the majority result in at least 3 phylogenetic signal detection methods (Supplemental File 1, table S2).

Thus, in summary, although it is possible to generate an artefactual phylogenetic signal using a randomly simulated trait, this scenario is exceptionally unlikely. Moreover, the strength of the phylogenetic signal observed among all simulated arbitrary variables is low (table 2; Supplemental File 1, table S1 and S2 and S3) and considerably weaker than that reported in rubisco kinetic traits (Bouvier et al., 2021). For example, when considering Pagel’s lambda, the metric used to correct for phylogenetic effects in our downstream phylogenetic least squares regression analysis (Bouvier et al., 2021), on average fewer than 1 in 1,000 of the re-shuffled data or the randomly simulated data produced a signal as strong or stronger then was observed for the real rubisco kinetic data (excluding *K*_O_ which had no phylogenetic signal in the analysis across all angiosperms) (Figure 3A and 3B; Supplemental File 1, table S3). This result corresponds with the p-values reported for rubisco kinetic traits in our original analysis and thus confirms the robustness of the phylogenetic signal detection methods utilized Bouvier et al., (2021). Thus, in contrast to the claim of Tcherkez and Farquhar (2021), the probability that randomly distributed data would produce a significant phylogenetic signal is 1%, and the probability that this randomly distributed data would produce a signal comparable or stronger to that observed for real rubisco kinetics is less than 0.1%. This raises questions about the method used by Tcherkez and Farquhar (2021) to compute their significant phylogenetic signal in their randomly simulated trait given that no formal description or raw data was provided.

**Figure 3.**
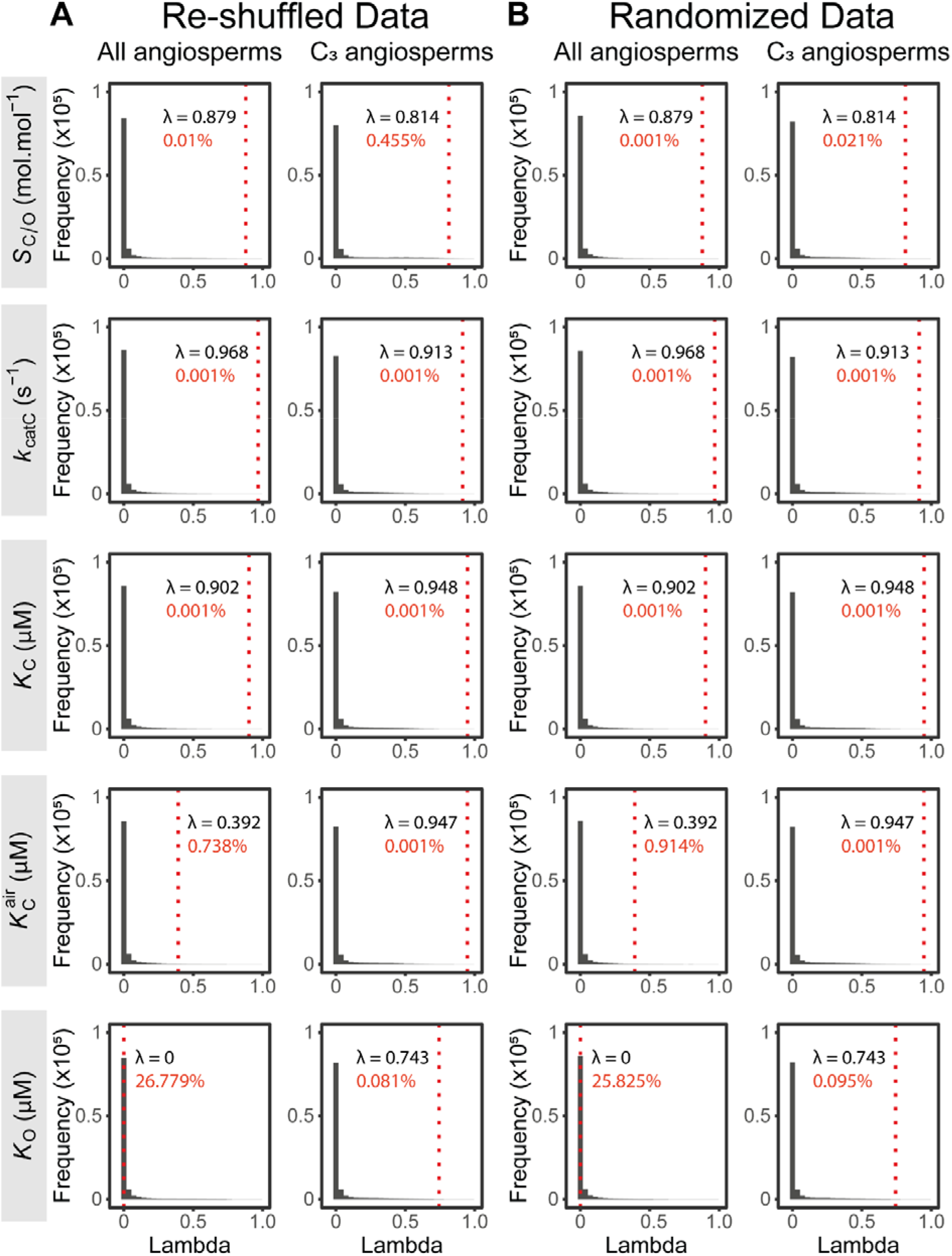
The phylogenetic signal strength in randomly simulated arbitrary trait data. A) Frequency distribution plots to show the phylogenetic signal assessed by Pagel’s lambda (λ) across 100,000 stochastically simulated datasets generated by randomly re-shuffling rubisco kinetic trait values across species on the phylogenetic tree. Each row represents data from a different rubisco kinetic trait. Data are shown for the analysis of all angiosperms and C_3_ angiosperms, respectively. The corresponding Pagel’s lambda value measured in the real rubisco kinetic trait data is included in each plot and is visualised by the red vertical dotted line. The percentage of arbitrary simulations which produced an equivalent or stronger phylogenetic signal than that observed for the real rubisco kinetic trait is also included in each plot. B) As in (A), but for 100,000 stochastically simulated trait datasets generated by randomly sampling from normal distributions inferred from the mean and standard deviation of each real rubisco kinetic trait. In both (A) and (B) it should be noted that the vast majority of randomly simulated arbitrary data produce a very weak phylogenetic signal which approaches 0, irrespective of the kinetic trait or the method of random arbitrary data simulation utilized. The average phylogenetic signal strength and percentage of significant tests in the randomly simulated arbitrary trait data based on Pagel’s lambda as well as the other signal detection methods utilized in the present study can be found in Supplemental File 1, table S1 and S2. The corresponding percentage of stochastically simulated datasets which produced an equivalent or stronger signal than that observed in the real rubisco kinetic traits, but inferred using the other phylogenetic signal detection methods can be found in Supplemental File 1, table S3.

### A phylogenetic signal is a universal feature of arbitrary traits distributed according to Brownian motion

In their Opinion article, Tcherkez and Farquhar (2021) further claimed that the phylogenetic signal in rubisco kinetics is artefactual because they can also observe the presence of a signal in a simulated trait that has been modelled to evolve on the rubisco tree by Brownian motion. However, this assertion demonstrates a misunderstanding about basic principles of phylogenetic data interrogation. To explain, in the context of comparative biology, Brownian motion is a widely used model of evolution in which a trait is simulated to evolve stochastically in both direction and magnitude through time. Specifically, in this model, traits evolve by accruing incremental changes along the branches of a phylogenetic tree (i.e., from the last common ancestor at the root to all extant species at the terminal tips) by sampling at each node from a random distribution of possible changes with zero mean and finite constant variance. In this way, Brownian motion is assumed to represent a tractable model to approximate how trait evolution might occur in the real world under a wide range of scenarios. This is because simulated variables acquire several inherent properties which parallel those typically observed during the evolution of many real biological traits. 1) Trait variation among extant species conforms to a normal distribution with a mean equal to the ancestral trait value (i.e., trait evolution is non-directional). 2) Trait variation among extant species increases as a function of evolutionary time encapsulated by the tree (i.e., trait evolution is additive). 3) Trait variation among extant species co-varies as a function of shared ancestry (i.e., traits are strongly phylogenetically structured). Given this latter property, it follows that phylogenetic signal will be a universal feature of all traits simulated on a tree using Brownian motion, irrespective of the underlying phylogenetic tree on which they are simulated. To illustrate this point, we have assessed the presence of phylogenetic signal in 100 arbitrary BM traits simulated to have evolved on each of 100,000 simulated trees and demonstrate that a strong significant phylogenetic signal is observed in ~100% of cases (Supplemental File 1, table S4). This result is recapitulated if the number of terminal tips on the simulated trees are set to equal either the full number of angiosperms in the rubisco dataset, or the reduced number which perform C_3_ photosynthesis (Supplemental File 1, table S4). In summary, therefore, the presence of phylogenetic signal in traits simulated by Brownian motion on a tree is ubiquitous and expected. Thus, the fact that Tcherkez and Farquhar (2021) found strong phylogenetic signal in an arbitrary trait that was guaranteed to have strong phylogenetic signal due to the manner in which it was simulated has no direct relevance to the analysis of phylogenetic signal in rubisco kinetic traits. It does not contradict the finding that rubisco kinetic traits have phylogenetic signal, nor does it negate the requirement to account for this phylogenetic signal in downstream statistical analysis of the kinetic trait data. The “serious statistical problem” in rubisco kinetics that exists is not caused by random effects in the data and needs to be correctly considered when computing trait correlations on a phylogenetic tree.

### Phylogenetic signal is not an artefact of biases in species sampling

Another criticism made by Tcherkez and Farquhar (2021) asserts that the presence of phylogenetic signal in rubisco kinetic traits is caused by biases in species sampling. Specifically, they argue that because groups of closely related species are present in the rubisco dataset, this has overestimated the phylogenetic signal in rubisco kinetic traits due to the effect of short branch lengths separating kinetically similar rubisco. However, as above, this argument demonstrates a fundamental confusion about the basic principles of phylogenetic data interrogation. For instance, phylogenetic signal detection methods work by computing the pairwise co-variation between trait similarity and phylogenetic distance across species. If nearly interchangeable rubisco kinetics are present in a dataset between closely related sister species, this would be accounted for in the phylogenetic context of their recent ancestry and short branch lengths, not overestimated by this effect (Wiens et al., 2008). In fact, the central premise of phylogenetic comparative methods is that they are robust to species sampling, whereas non-phylogenetic methods are not. To illustrate this point, we have removed the species hypothesized to be problematic by Tcherkez and Farquhar (2021) from the analysis of all angiosperms and C_3_ angiosperms (including plants in the *Oryza* clade, as well as those belonging to both the *Aegilops* and closely related *Triticum* clades which together share a convoluted evolutionary history (Petersen et al., 2006)) and have shown that an analogous phylogenetic signal is observed in this analysis (Supplemental File 1, table S5) as compared to in the original analysis when these species were present (table 2). This conclusion is true whether all angiosperms, or whether only C_3_ angiosperms are considered (Supplemental File 1, table S5; table 2). In summary, therefore, biases in species sampling including short branches separating nearly identical sister species, and overrepresentation of certain groups, are not responsible for the phylogenetic signal in rubisco kinetic traits. Indeed, it has in fact been shown that short branches have the inverse effect of causing phylogenetic signal to be underestimated due to complications arising with difficulties in resolving the correct phylogenetic history of species (Wiens et al., 2008).

### Phylogenetic signal in rubisco kinetics is not an artefact of using rbcL-based trees

A further issue raised by Tcherkez and Farquhar (2021) is associated with the use of the *rbcL* gene to reconstruct the phylogenetic tree of species in our analysis. Specifically, they contend that tree inference based on *rbcL* sequence similarity across species may have caused inflation of phylogenetic signal due to overfitting. This is because mutations in *rbcL* which contribute to kinetic variation among species would also affect tree topology. However, we specifically addressed this possibility in our original study (Bouvier et al., 2021) and have previously shown this to not be an issue as we were able to replicate our results when using a phylogeny generated from only *rbcL* codon positions which contained synonymous substitutions across species (i.e., where columns in the alignment containing non-synonymous substitutions that could affect enzyme kinetics were removed) (Bouvier et al., 2021). Given that the use of this tree did not impact our analysis compared to that based on the complete *rbcL* tree, the phylogenetic signal in rubisco kinetics was confirmed to not be attributable to overfitting (Bouvier et al., 2021). As such, the full alignment *rbcL* tree was deemed appropriate for our analyses given that it has a long history in phylogenetic inference of species relationships (APG, 2016, 1998; Gielly and Taberlet, 1994) and more accurately reflects the evolutionary history of rubisco compared to the partial *rbcL* tree inferred from only synonymous substitutions.

Owing to limited availability of publicly sequenced nuclear (*n* = 26) or chloroplast (*n* = 8) genomes for species in the kinetic dataset, it was not possible to use whole genome phylogenomic approaches to infer the tree of species in our analysis. Nevertheless, to provide further reassurance in the present study that the phylogenetic signal in rubisco kinetics is not a consequence of the use of *rbcL* for phylogenetic inference, we repeated our analysis using phylogenies inferred from the coding sequences of several other chloroplast encoded genes which are frequently utilized in systematics for species classification (Bohs and Olmstead, 1997; Ferguson, 1998; Hilu and Liang, 1997; Koch et al., 2001; Savolainen et al., 2000; Wolf, 1997; Yen and Olmstead, 2000) and importantly are unrelated to the kinetic traits of interest. Specifically, genes employed to infer these respective alternative trees include the ATP synthase beta subunit (*atpB*), the NADH dehydrogenase F (*ndhF*) subunit and maturaseK (*matK*) which was specifically argued by Tcherkez and Farquhar (2021) to reflect the evolutionary relationships of the C_4_ species more accurately in our analysis compared to those inferred by *rbcL*. Analogous measurements of phylogenetic signal to those reported in our original paper are observed when these alternative gene trees are used (Supplemental File 1, table S6; table 2). In summary, therefore, the presence of phylogenetic signal in rubisco kinetics is not an artefact of using *rbcL*-based trees to infer phylogeny.

### Phylogenetic signal is not an artefact of method-to-method variability in trait measurements

A separate concern raised by Tcherkez and Farquhar (2021) is related to the fact that our kinetic data was compiled from numerous sources in the literature. Specifically, in the analysis of angiosperms, the kinetic dataset was compiled across ten independent studies, including (Galmés et al., 2014; Kubien et al., 2008; Occhialini et al., 2016; Orr et al., 2016; Prins et al., 2016; Savir et al., 2010; Sharwood et al., 2016; von Caemmerer et al., 1994; Whitney et al., 2011; Zhu et al., 1998). Given that laboratory-to-laboratory variability exists in kinetic measurements due to inconsistencies in experimental protocols and techniques (Iñiguez et al., 2021), it was argued that such biases could confound the results of a phylogenetic signal analysis. Theoretically, this effect would be further exacerbated if studies have characterized rubisco among closely related clusters of species on the tree of life, which would thus have more comparable values to one another, but less comparable values to other distantly related species characterized in separate laboratories.

Although we agree that methodological-induced systematic biases are a genuine concern in analysis of any metadata, we have previously taken every effort to mitigate these effects from influencing our analysis. For example, the rubisco meta-dataset originally assembled by Flamholz et al., (2019) was limited to include only measurements assayed under conditions of 25 °C and pH 7.8–8.0. Moreover, rubisco *S*_C/O_ values in Bouvier et al., (2021) were normalised between studies using an oxygen electrode assay (Parry et al., 1989) and those using the improved high precision gas-phase-controlled ^3^H-RuBP-fixation assay (Kane et al., 1994) by applying corrections based on wheat as an internal standard in both methods. Finally, study-specific errors were also limited in Bouvier et al., (2021) by taking averages across rubisco kinetics in the same species which were measured independently by different authors. Therefore, although it is impossible to completely avoid measurement variability across published studies, we have been pragmatic to lessen this impact where possible by adopting caution in data compilation (Flamholz et al., 2019) as well as in subsequent data processing (Bouvier et al., 2021).

Nevertheless, to confirm the robustness of our results to inter-study biases in measurements, we have repeated our analysis here using data derived from a single study in the meta-dataset so that there can be no influence of method-to-method or laboratory-to-laboratory variability. For this purpose, the subset of rubisco characterized in Orr et al., (2016) were analyzed, as it had sufficient species sampling for a robust statistical analysis. Phylogenetic interrogation of this dataset revealed a similar magnitude and significance of phylogenetic signal in rubisco (Supplemental File 1, table S7) compared to that which was originally reported from the meta-dataset (table 2). This same outcome is observed irrespective of whether all angiosperms, or exclusively C_3_ angiosperms, were analysed (Supplemental File 1, table S7; table 2). Therefore, phylogenetic signal in rubisco kinetics is not an artefact of method-to-method or laboratory-to-laboratory variability in trait measurements.

In addition to the general concerns with our meta-analysis that are addressed above, a more specific criticism of method-to-method biases in our dataset was associated with the *Limonium* tribe. In particular, Tcherkez and Farquhar (2021) argue that the *S*_C/O_ values of *Limonium* originally measured by (Galmés et al., 2014) as found in our dataset have been overestimated by ~13 mol mol^-1^ as compared to all other measured rubisco. Accordingly, they have claimed that this bias was the cause of the phylogenetic signal in this trait in our study. However, in the present analysis when such a correction of *S*_C/O_ values in *Limonium* is carried out, we find that this does not cause an appreciable difference in the magnitude of the phylogenetic signal in this trait either across all angiosperms, or across C_3_ angiosperms (Supplemental File 1, table S8), as compared to the original analyses based on uncorrected values (table 2). An analogous conclusion is similarly obtained across both the analysis of all angiosperms and C_3_ angiosperms if species in the *Limonium* clade were removed completely (Supplemental File 1, table S9; table 2). In combination, therefore, phylogenetic signal in rubisco kinetic traits is genuine and is not an artefact of the sampled species nor the data being compiled from different sources.

### Phylogenetic signal is not an artefact of C_3_ vs C_4_ photosynthesis

Given our analysis is robust to all criticisms made by Tcherkez and Farquhar (2021), an interesting question of fundamental importance is why strong pervasive phylogenetic signal exists in rubisco kinetics (Bouvier et al., 2021). On this matter, despite their objections, Tcherkez and Farquhar (2021) postulate that this phenomenon is primarily driven by differences in the biochemical niche of the enzyme across the tree of life. Specifically, given the widespread kinetic differences associated with adaptation to intracellular CO_2_ levels in plants with C_3_ (higher CO_2_ specificity) and C_4_ metabolisms (higher CO_2_ turnover) (Bouvier et al., 2021), Tcherkez and Farquhar (2021) claim that C_3_ vs. C_4_ kinetic differences are responsible for the observed phylogenetic signal.

However, the above hypothesis put forward by Tcherkez and Farquhar (2021) is flawed by confusion between global phylogenetic signal (referred to here and in Bouvier et al., (2021) as simply “phylogenetic signal”) and local phylogenetic signal (analysed in Tcherkez and Farquhar, (2021), but not discussed in Bouvier et al., (2021) or in the present study until this point). In brief, local measures of phylogenetic signal, such as local Moran’s *I* (Anselin, 1995), serve to identify unique and interesting patterns of trait variation in a group of species which are not conserved in neighbouring clades (i.e., the kinetic outliers of rubisco in C_4_ compared to C_3_ sister species along the angiosperm tree). In contrast, global phylogenetic signal as assessed here and by Bouvier et al., (2021) is different and instead characterises the average statistical co-variation between trait values and phylogenetic distances across all pairwise combinations of species on the tree. As such, global metrics of phylogenetic signal (including Pagel’s lambda (Pagel, 1999), Blomberg’s *K* and *K*\* (Blomberg et al., 2003), Moran’s *I* (Gittleman and Kot, 1990) and Abouheif’s *Cmean* (Abouheif, 1999)) are ultimately used to correct for the phylogenetic non-independence of data in phylogenetic comparative models. Based on this distinction, although kinetic changes associated with transition to C_4_ photosynthesis in discrete pockets on the phylogenetic tree will produce hotspots of local phylogenetic signal, this effect will reduce the estimated global phylogenetic signal which we measure. This is because the convergence and homoplasy of the C_4_ trait weakens the average statistical dependence between trait similarity and phylogenetic relatedness among the group of sampled species as a whole (Hansen and Martins, 1996; Kamilar and Cooper, 2013). Indeed, this fact was shown in the results of our original study (Bouvier et al., 2021) as well as those presented here (table 2; Supplemental File 1, table S4–S9) whereby inclusion of C_4_ and other C_3_-C_4_ intermediary species reduces the signal strength relative to when only C_3_ angiosperms are considered. Moreover, this result is also observed in the analysis presented by Tcherkez and Farquhar (2021), although this effect is misinterpreted by these authors. Specifically, using *k*_catC_ as an example Tcherkez and Farquhar (2021) show that C_4_ angiosperms are associated with local phylogenetic signal when all species are considered, however they detect no global phylogenetic signal in this analysis (Figure 2 of Tcherkez and Farquhar, (2021)). In contrast, when C_4_ and other intermediary C_3_-C_4_ and C_4_-like species are ignored, Tcherkez and Farquhar (2021) show the local phylogenetic signal associated with these groups is lost and global phylogenetic signal accordingly becomes significant at short phylogenetic distances of the remaining species (Figure 2 of Tcherkez and Farquhar, (2021)). This result demonstrates that although C_3_ vs. C_4_ kinetic differences drive local phylogenetic signal, they are not responsible (and in fact weaken) the global phylogenetic signal which is of importance to the discussion of Bouvier et al., (2021). This was already discussed in our original study (Bouvier et al., 2021). Thus, the contention of Tcherkez and Farquhar (2021) that C_4_-mediated kinetic changes are responsible for the observed phylogenetic signal across the whole tree is nonsensical, given that this signal is detected when analysing only C_3_ angiosperms (as originally reported (Bouvier et al., 2021)), and the strength of the signal in the analysis of C_3_ angiosperms is greater than when both C_3_ and C_4_ angiosperms are considered together.

## Discussion

Rubisco (ribulose-1,5-bisphosphate [RuBP] carboxylase/oxygenase) is the primary entry point for carbon into the biosphere (Field et al., 1998; Tabita et al., 2008) and accordingly, is the source of almost all organic carbon which has ever existed. As such, gaining an appreciation of how rubisco has evolved is of fundamental importance to our understanding of life on our planet. Interestingly, however, rubisco has long presented an evolutionary paradox. This is because, despite the central importance of rubisco in underpinning host autotroph metabolism and ultimately the global food chain, the enzyme is considered by many to be an inefficient catalyst under the high CO_2_, low O_2_ conditions of the present day. This is due to the fact that rubisco has a modest turnover rate for its primary CO_2_ reaction (Badger et al., 1998; Bar-Even et al., 2011) and also catalyses a costly secondary reaction with O_2_ that culminates in the loss of previously fixed carbon (Bowes et al., 1971; Chollet, 1977; Sharkey, 2020). Although an updated examination has suggested that rubisco’s kinetics are overall perhaps not as bad as often assumed (Bathellier et al., 2018), these puzzling catalytic limitations of the enzyme have nevertheless caused rubisco to be an enigma and the source of intensive and sustained research interest for over 40 years.

Until our phylogenetically resolved investigation of rubisco kinetic evolution (Bouvier et al., 2021), the principal hypothesis forwarded to explain the above “rubisco paradox” proposed that severe catalytic trade-offs exist between the kinetic traits of the enzyme. Specifically, this theory was first pioneered by two cross-species analyses (Savir et al., 2010; Tcherkez et al., 2006) which reported strong antagonistic correlations between rubisco kinetic traits and posited that these correlations were caused by unavoidable chemical constraints on its catalytic mechanism (despite a limited understanding of this chemical mechanism). In this way, these kinetic trait interdependencies were understood to provide a ceiling on rubisco optimisation by limiting the adaptative capacity of its individual kinetic traits (Savir et al., 2010; Tcherkez et al., 2006). However, both these analyses which first described the rubisco kinetic trait trade-offs (Savir et al., 2010; Tcherkez et al., 2006), in addition to all subsequent analyses which have re-investigated these trade-offs among different taxa (Flamholz et al., 2019; Iñiguez et al., 2020; Young et al., 2016), failed to account for the phylogenetic context of the sampled species being considered in the analysis. This is a serious statistical problem because species (and the rubisco they encode) cannot be treated as independent observations in comparative studies owing to the shared ancestry of all organisms on the hierarchical tree of life (Felsenstein, 1985). Thus, the failure to account for phylogenetic non-independence of rubisco data in previous studies has violated one of the core assumptions of the conventional statistical methods which were applied (i.e., independence of residuals; table 1; Figure 1) meaning that the results drawn from these analyses may be, wholly or in part, artefactual.

To assess whether the methodological flaws of previous cross-species analyses has led to mistaken inferences about rubisco, in Bouvier et al., (2021) we re-investigated the kinetic evolution of this enzyme in a phylogenetically resolved manner. We discovered for the first time that rubisco kinetic traits exhibit strong and significant phylogenetic signal (Bouvier et al., 2021). The presence of this signal means that kinetic trait values are statistically similar across enzymes as a function of their shared evolutionary history (i.e., the kinetic traits exhibit strong phylogenetic non-independence). Accordingly, we re-evaluated the correlations between rubisco kinetic traits when accounting for this phylogenetic signal and found that only canonical trade-offs between the Michaelis constant for CO_2_ (*K*_C_) and carboxylase turnover (*<u>k</u>*_catC_), and between the Michaelis constants for CO_2_ and O_2_ (*K*_O_) were robust to phylogenetic effects (Figure 2A – 2C) (Bouvier et al., 2021). Though, the strength of these trade-offs were weak and considerably attenuated relative to earlier estimates (Figure 2A – 2C) (Bouvier et al., 2021). Therefore, although catalytic trade-offs exist, these are not absolute, and have been previously overestimated due to phylogenetic biases in the kinetic data (Figure 2A – 2C). Instead, kinetic traits have been able to evolve largely independently of one another during the diversification of rubisco. Moreover, when we further investigated the constraints on the enzyme, we found that phylogenetic constraints explained more variation in rubisco kinetics, and have thus had a larger impact on limiting enzyme adaptation, compared to the combined action of all catalytic trade-offs (Bouvier et al., 2021). In summary, therefore, although rubisco catalytic trade-offs exist as determined by chemistry, these represent a less serious constraint on rubisco kinetic adaptation compared to previous assumptions. Instead, phylogenetic constraints, likely caused by slow molecular evolution in rubisco (Bouvier et al., 2022) and more general constraints on the molecular evolution of chloroplast encoded genes (Robbins and Kelly, 2022), have presented a more significant barrier to improved rubisco catalytic efficiency.

In Bouvier et al., (2021) we made every effort to carefully describe the method by which we have computed our phylogenetic signal in rubisco kinetic data. In addition, we were also conscious to discuss what this phylogenetic signal means, as well as to explain in detail how the presence of this signal has incorrectly resulted in overestimated kinetic trait correlation coefficients in previous analyses. However, our results have been brought under criticism by Tcherkez and Farquhar, (2021). Specifically, these authors have argued that the phylogenetic signal we detect in rubisco kinetics is a consequence of computational artefacts due to a combination of biases associated with species sampling (including the use of near identical sister species as well as the overrepresentation of certain groups), the use of *rbcL-based* trees for phylogenetic inference, method-to-method and laboratory-to-laboratory variability among kinetic measurements in the rubisco meta-dataset, and finally, homoplasy of the C_4_ trait across sampled species. On this matter, Tcherkez and Farquhar (2021) argue that our phylogenetic signal in rubisco kinetics is not valid because they can also generate a signal using our method with both a randomly simulated arbitrary trait as well as a trait distributed according to Brownian motion. In the present response, we critically review each of these claims on a point-by-point basis by re-visiting and extending our original analysis. In doing so, we provide unequivocal evidence that all of the criticisms raised by Tcherkez and Farquhar (2021) were either incorrect, arose from a fundamental misunderstanding of basic phylogenetic concepts, or were already addressed in our original manuscript and proven to be unimportant. As such, we reaffirm that our results in Bouvier et al., (2021) are robust. Thus, our original conclusions, as summarized above, are unaltered.

To avoid any confusion on the points made by Tcherkez and Farquhar, (2021), it is worthwhile discussing further the subject of C_3_ vs. C_4_ photosynthesis in the context of measuring phylogenetic signal in rubisco kinetics. As described by our previous results (Bouvier et al., 2021) and those in (Capó-Bauçà et al., 2022; Cummins, 2021; Iñiguez et al., 2020), we agree that kinetic adaptations in rubisco occur in conjunction with the emergence of CO_2_-concentrating mechanisms. For example, rubisco evolve to become faster (increased carboxylase turnover) and less specific (decreased specificity and CO_2_ affinity) during transition from C_3_ to C_4_ photosynthesis (Bouvier et al., 2021). Nevertheless, although co-evolution between rubisco and C_4_ photosynthesis is apparent, as underpinned by changes in the gaseous micro-environment of the enzyme, this is not responsible for the phylogenetic signal in rubisco kinetic traits as proposed by Tcherkez and Farquhar (2021). Indeed, rubisco kinetic adaptation associated with the convergent evolution of C_4_ photosynthesis in discrete clusters of species on the phylogenetic tree in fact has the opposite effect of degrading the pairwise statistical association between phylogenetic distance and kinetic distance across all sampled rubisco as a whole. Thus, although differences in prevailing gaseous conditions around rubisco exist between C_3_ and C_4_ species and underpin differences in the trajectory of rubisco adaptation, this does not drive (and in fact weakens) the observed phylogenetic signal in rubisco kinetics.

Although the present work is focussed on angiosperms, it should be noted that we have previously observed identical results in rubisco across the tree of life (table 2) (Bouvier et al., 2021). Thus, rubisco evolution is only weakly constrained by catalytic trade-offs and is instead more limited by phylogenetic constraint. We propose that this phylogenetic constraint arises from a combination of a high degree of purifying selection (Robbins and Kelly, 2022), the requirement for high levels of transcript and protein abundance (Kelly, 2018; Robbins and Kelly, 2022; Seward and Kelly, 2018), the requirement for maintaining complementarity to a wide array of molecular chaperones which assist in protein folding and assembly (e.g., Raf1, Raf2, RbcX, BSD2, Cpn60/Cpn20) and metabolic regulation (e.g., rubisco activase) (Aigner et al., 2017; Carmo-Silva et al., 2015), and finally, the need to preserve overall protein stability within the molecular activity-stability trade-offs (Cummins et al., 2018; Duraõ et al., 2015; Studer et al., 2014). These factors combined contribute to the exceedingly slow rate of molecular evolution in *rbcL* (Bouvier et al., 2022) which presents a major barrier on rubisco optimisation.

## Conclusions

Prior to our phylogenetically resolved investigation of rubisco evolution (Bouvier et al., 2021), all comparative studies which have measured correlations between rubisco kinetic traits were computed in the absence of accounting for the phylogenetic non-independence of the data. We described how this omission has led to the mistaken inference that the biochemical landscape of rubisco is severely constrained by catalytic trade-offs (Bouvier et al., 2021). In the present study, we revisit these analyses to address a series of criticisms of our work that were put forward in a recent opinion article (Tcherkez and Farquhar, 2021). Here, we demonstrate that all of these criticisms were either misguided or incorrect, and thus our original conclusions are unaltered. Namely, strong phylogenetic signal exists in rubisco kinetic traits and cause an overestimation of catalytic trade-offs unless correctly accounted for. In actual fact, phylogenetic constraints have limited rubisco kinetic adaptation to a greater extent than the combined action of catalytic trade-offs. These phylogenetic constraints are caused by multiple evolutionary factors act to limit rubisco molecular sequence evolution (Bouvier et al., 2022) and combined, help to explain why the enzyme is poorly efficient under present-day conditions but better adapted to the former high CO_2_, low O_2_ atmosphere in which it evolved. These conclusions agree with recent studies which have also emphasized that kinetic trait correlations are generally too weak to support the rubisco trade-off model (Cummins et al., 2019, 2018; Flamholz et al., 2019; Galmés et al., 2014) as well as experimental results from rubisco engineering efforts which have been able to successfully produce enzyme variants that deviate from proposed catalytic trade-offs (Wilson et al., 2018; Zhou and Whitney, 2019).

## Materials and Methods

### Kinetic data

The rubisco kinetic dataset used in this work was compiled across various sources in the primary literature (Bouvier et al., 2021; Flamholz et al., 2019). This data includes experimentally determined measurements of wild-type rubisco assayed under conditions of pH 7.8–8.0 and 25 °C for CO_2_/O_2_ specificity (*S*_C/O_), maximum carboxylase turnover rate per active site (*k*_catC_), and the respective Michaelis constant (i.e., the substrate concentration at half saturated catalysed rate) for both CO_2_ (*K*_C_) and O_2_ (*K*_O_) substrates. In addition, measurements of the Michaelis constant for CO_2_ in 20.95% O_2_ ambient air (*K*_C_^air^) were also available as previously derived from *K*_C_ and *K*_O_ using the formula *K*_C_ + (*K*_C_ [O_2_]/*K*_O_), where 20.95% [O_2_] in water is 253□μM (Bouvier et al., 2021). As previously described (Bouvier et al., 2021), modifications of this dataset have been performed by normalizing between *S*_C/O_ measurements that were determined using an oxygen electrode assay (Parry et al., 1989) and those determined using the high precision gas-phase-controlled ^3^H-RuBP-fixation assay (Kane et al., 1994) by using wheat as an internal standard in both. In addition, this dataset was also previously modified to average across duplicate entries of species (including synonyms) between studies (Bouvier et al., 2021). It should be noted that in the present study, only data for the 137 angiosperm species with complete measurements of *S*_C/O_, *k*_catC_, *K*_C_, *K*_C_^air^ and *K*_O_ were considered given that this is the taxonomic group which was solely focused on by Tcherkez and Farquhar (2021). Further, as before, all kinetic traits were log transformed prior to interrogation to satisfy the distributional assumptions of the statistical tests being performed.

### Phylogenetic tree inference based on rbcL and other gene sequences

All phylogenetic trees inferred from the coding sequence of the rubisco large subunit (*rbcL*) were obtained from our previous work (Bouvier et al., 2021). This includes the separate *rbcL* trees which describes the evolutionary history of all angiosperms, as well as the subset of angiosperms which perform C_3_ photosynthesis. Phylogenetic trees based on other chloroplast encoded genes including maturaseK (*matK*), ATP synthase beta subunit (*atpB*), and the NADH dehydrogenase F (*ndhF*) subunit were generated in the present study following the exact method previously described (Bouvier et al., 2021). Specifically, coding sequences of *matK, atpB*, and *ndhF* were obtained from NCBI (https://www.ncbi.nlm.nih.gov/) for as many of the angiosperm species in the kinetic dataset as possible. Multiple sequence alignments were generated for each respective gene using mafft L-INS-i (Katoh and Standley, 2013), and alignments were processed to remove any partial, chimeric, or erroneously annotated sequences. Finally, bootstrapped maximum-likelihood phylogenetic trees were inferred from each non-gapped sequence alignment by IQ-TREE (Nguyen et al., 2015) using the ultrafast bootstrapping method with 1,000 replicates and the Shimodaira–Hasegawa approximate-likelihood ratio branch test, with the best fitting model of sequence evolution chosen automatically. All trees were rooted manually using Dendroscope (Huson and Scornavacca, 2012). For each of these inferred trees based on *matK*, *atpB*, and *ndhF* gene sequences, a corresponding tree containing only C_3_ species was obtained using the function drop.tip from the ‘ape’ package (Paradis et al., 2004; Paradis and Schliep, 2019) in the R environment.

Due to the differences in phylogenetic information available for tree building from the alignments of *matK*, *atpB*, and *ndhF* genes as compared to the alignment based on the *rbcL* gene, additional polytomies which include species with terminal zero-length branches were detected in these alternative chloroplast gene trees (due to species possessing 100% sequence identity for the gene in question). However, it was not appropriate to condense these zero-branch length nodes into single datapoints as previously performed in the analysis based on the *rbcL*-based trees (Bouvier et al., 2021). This is because these other chloroplast genes are unrelated to rubisco kinetic traits. Thus, averaging kinetic data across zero-branch length species on these other trees is incorrect in the context of the downstream phylogenetic analysis being performed, given that these nodes are known to possess differences in their rubisco sequences which are not captured by the tree. Thus, to get around this issue and maximize the number of species taken forward for analysis, zero-terminal length branches in *matK*, *atpB*, and *ndhF* trees were arbitrarily resolved into fully bifurcating trees by using the function multi2di in the ‘ape’ R package (Paradis et al., 2004; Paradis and Schliep, 2019) and increasing terminal branch length values by a minimal value of 0.000001. The total number of angiosperms and the total number of C_3_ angiosperms encapsulated by each of these alternative gene trees can be found in (Supplemental File 1, table S6). These alternative trees are provided in Newick format in (Supplemental File 2).

### Generation of randomly simulated arbitrary traits

To evaluate whether randomly simulated data could produce an artefactual phylogenetic signal, two separate approaches were employed. First, 100,000 random permutations of the data were performed manually such that in each permutation the kinetic trait data for the species set was subject to a Fisher-Yates shuffle. Each permutation was subject to phylogenetic signal analysis as described below and the proportion of permutations that obtained an equivalent or larger phylogenetic signal to that observed in the real kinetic trait data were recorded. Second, 100,000 stochastically simulated trait datasets were generated by randomly sampling from normal distributions inferred from the mean and standard deviation of each real rubisco kinetic trait. To achieve this, the rnorm function was employed from the ‘compositions’ package (van den Boogaart and Tolosana-Delgado, 2008) in the R environment. Each simulation was subject to phylogenetic signal analysis as described below and the proportion of simulated trait dataset that obtained an equivalent or larger phylogenetic signal to that observed in the real kinetic data were recorded.

### Simulation of kinetic traits on phylogenetic trees by Brownian motion

To show that a phylogenetic signal is a universal feature of all traits which are distributed according to Brownian motion on any underlying phylogenetic tree, 100,000 bifurcating trees were randomly simulated using the rtree function from the ‘ape’ package (Paradis et al., 2004; Paradis and Schliep, 2019) in R. This process was repeated for trees containing both the same number of terminal tips as the full set of angiosperms in the rubisco kinetic dataset and the same number of tips as the subset of C_3_ angiosperms, respectively. Next, for each tree, 100 quantitative traits were simulated to evolve under a standard Brownian motion model using the function fastBM in the R package ‘phytools’ (Revell, 2012). In total, this resulted in 10,000,000 unique Brownian motion trait simulations for the analysis of all angiosperms and C_3_ angiosperms, respectively. Each simulated trait was subject to phylogenetic signal analysis using the methods outlined below.

### Phylogenetic signal analysis

The magnitude and significance of phylogenetic signal in a given trait on a given underlying phylogenetic tree was computed using five independent detection methods, including Pagel’s lambda (Pagel, 1999), Moran’s I (Gittleman and Kot, 1990), Abouheif’s Cmean (Abouheif, 1999) and Blomberg’s K and K* (Blomberg et al., 2003). Specifically, these tests work by assessing the statistical association between trait and phylogenetic similarity among all pairwise combinations of species on the tree. Implementation of these phylogenetic signal detection tests was performed in the R environment as previously described (Bouvier et al., 2021).

## Acknowledgements

This work was funded by the Royal Society. JWB was funded by the BBSRC through BB/J014427/1.

## Data Availability

All data used in this study is provided in the supplemental material.

## Author Contributions

JWB and SK performed the computational analysis and wrote the manuscript.

## Boxes

### Box 1.

A list of definitions for key terms as they are applied in the context of the present study and in our original study (Bouvier et al., 2021).

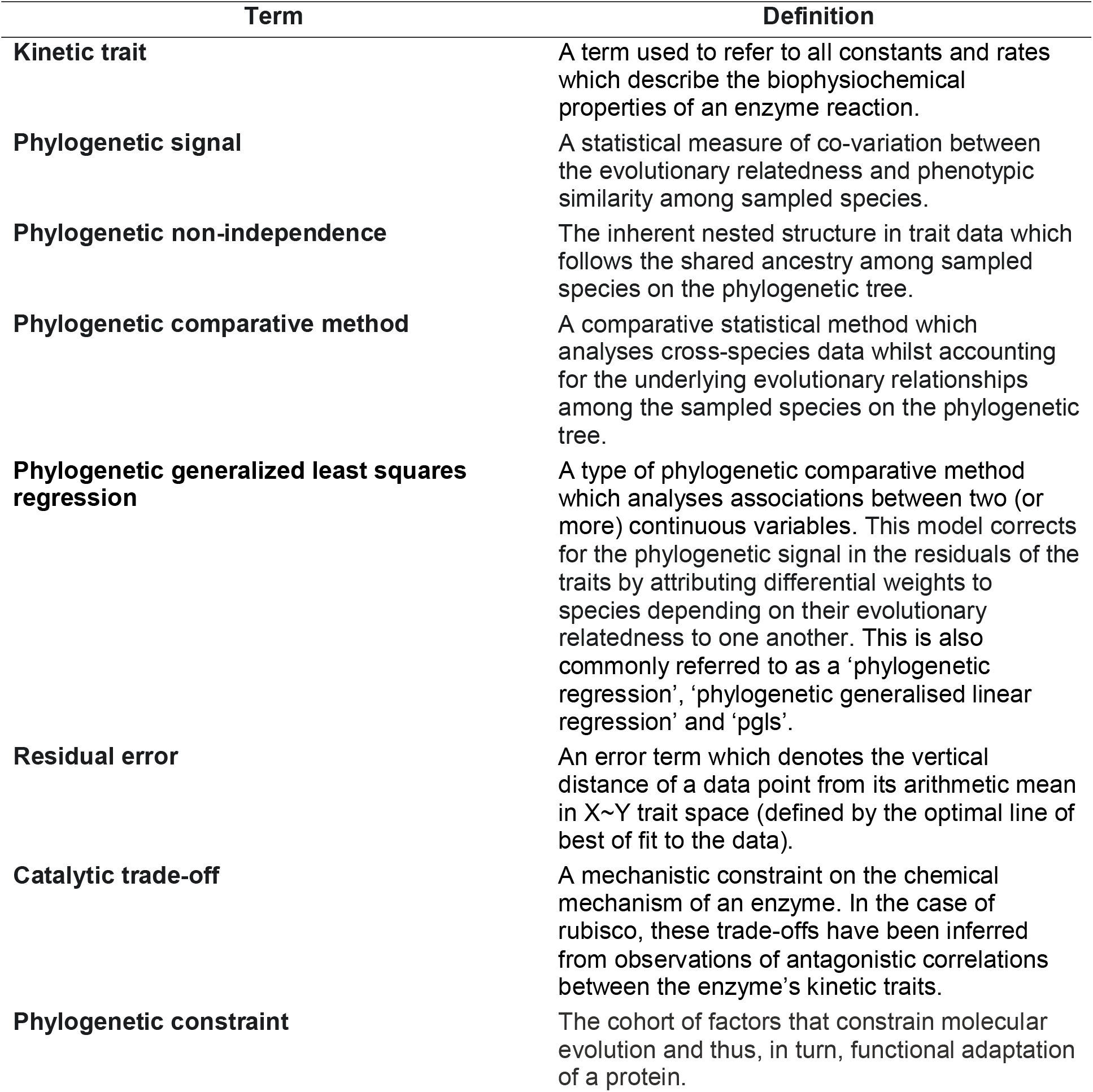

## Notes

### Competing Interest Statement

The authors have declared no competing interest.

