## Supplemental File 1 for "Response to Tcherkez and Farquhar: Rubisco adaptation is more limited by phylogenetic constraint than by catalytic trade-off"

### Supplemental Tables

**Table S1.** The phylogenetic signal strength (mean ± S.E.) and percent of significant tests in arbitrary trait data generated by randomly re-shuffling real rubisco kinetic trait data across species on the phylogenetic tree using 100,000 replicates. Phylogenetic signal is measured using five signal detection methods. Data are shown from the analysis of all angiosperms and C_3_ angiosperms, respectively.

| **Data** | **Arbitrary Trait** | ***Cmean*** | | ***I*** | | ***K*** | | ***K**** | | ***Lambda*** | | **Consensus** |
| --- | --- | --- | --- | --- | --- | --- | --- | --- | --- | --- | --- | --- |
|  |  | ***Stat*** | ***% Sig*** | ***Stat*** | ***% Sig*** | ***Stat*** | ***% Sig*** | ***Stat*** | ***% Sig*** | ***Stat*** | ***% Sig*** | ***% Sig*** |
| **All Angiosperms** | ***S*_C/O_** | -0.008 ± 0 | 5.052 | -0.008 ± 0 | 5.059 | 0 ± 0 | 5.165 | 0 ± 0 | 5.133 | 0.031 ± 0 | 1.817 | 1.075 |
|  | ***k*_catC_** | -0.009 ± 0 | 5.019 | -0.008 ± 0 | 4.994 | 0 ± 0 | 4.963 | 0 ± 0 | 5.066 | 0.017 ± 0 | 1.132 | 1.656 |
|  | ***K*_C_** | -0.008 ± 0 | 5.080 | -0.009 ± 0 | 4.898 | 0 ± 0 | 5.015 | 0 ± 0 | 4.92 | 0.018 ± 0 | 1.163 | 1.448 |
|  | ***K*_C_^air^** | -0.009 ± 0 | 4.882 | -0.009 ± 0 | 4.951 | 0 ± 0 | 5.059 | 0 ± 0 | 5.097 | 0.018 ± 0 | 1.174 | 1.473 |
|  | ***K*_O_** | -0.009 ± 0 | 4.945 | -0.009 ± 0 | 4.902 | 0 ± 0 | 4.898 | 0 ± 0 | 4.946 | 0.026 ± 0 | 1.464 | 1.061 |
| **C_3_ Angiosperms** | ***S*_C/O_** | -0.012 ± 0 | 5.000 | -0.011 ± 0 | 5.086 | 0.001 ± 0 | 5.097 | 0.001 ± 0 | 5.053 | 0.056 ± 0 | 3.055 | 1.242 |
|  | ***k*_catC_** | -0.012 ± 0 | 4.964 | -0.011 ± 0 | 5.043 | 0 ± 0 | 5.023 | 0 ± 0 | 4.954 | 0.032 ± 0 | 1.419 | 1.625 |
|  | ***K*_C_** | -0.012 ± 0 | 4.997 | -0.012 ± 0 | 4.951 | 0 ± 0 | 5.054 | 0 ± 0 | 5.049 | 0.034 ± 0 | 1.521 | 1.460 |
|  | ***K*_C_^air^** | -0.011 ± 0 | 5.014 | -0.011 ± 0 | 4.867 | 0 ± 0 | 5.013 | 0 ± 0 | 5.048 | 0.034 ± 0 | 1.596 | 1.439 |
|  | ***K*_O_** | -0.011 ± 0 | 4.992 | -0.011 ± 0 | 5.059 | 0 ± 0 | 5.018 | 0 ± 0 | 5.015 | 0.036 ± 0 | 1.671 | 1.485 |

**Table S2.** The phylogenetic signal strength (mean ± S.E.) and percent of significant tests in arbitrary trait data generated by randomly simulating data with the same distributional properties as each real rubisco kinetic trait using 100,000 replicates. Phylogenetic signal is measured using five signal detection methods. Data are shown from the analysis of all angiosperms and C_3_ angiosperms, respectively.

| **Data** | **Arbitrary Trait** | ***Cmean*** | | ***I*** | | ***K*** | | ***K**** | | ***Lambda*** | | **Consensus** |
| --- | --- | --- | --- | --- | --- | --- | --- | --- | --- | --- | --- | --- |
|  |  | ***Stat*** | ***% Sig*** | ***Stat*** | ***% Sig*** | ***Stat*** | ***% Sig*** | ***Stat*** | ***% Sig*** | ***Stat*** | ***% Sig*** | ***% Sig*** |
| **All Angiosperms** | ***S*_C/O_** | -0.009 ± 0 | 4.994 | -0.008 ± 0 | 5.101 | 0 ± 0 | 4.912 | 0 ± 0 | 4.957 | 0.018 ± 0 | 1.232 | 1.515 |
|  | ***k*_catC_** | -0.009 ± 0 | 5.050 | -0.008 ± 0 | 4.931 | 0 ± 0 | 4.947 | 0 ± 0 | 4.915 | 0.018 ± 0 | 1.208 | 1.444 |
|  | ***K*_C_** | -0.008 ± 0 | 5.100 | -0.008 ± 0 | 5.160 | 0 ± 0 | 4.966 | 0 ± 0 | 5.040 | 0.019 ± 0 | 1.201 | 1.569 |
|  | ***K*_C_^air^** | -0.008 ± 0 | 5.098 | -0.008 ± 0 | 5.117 | 0 ± 0 | 5.087 | 0 ± 0 | 5.114 | 0.018 ± 0 | 1.260 | 1.547 |
|  | ***K*_O_** | -0.009 ± 0 | 4.941 | -0.009 ± 0 | 4.904 | 0 ± 0 | 4.990 | 0 ± 0 | 4.973 | 0.018 ± 0 | 1.192 | 1.434 |
| **C_3_ Angiosperms** | ***S*_C/O_** | -0.012 ± 0 | 4.957 | -0.012 ± 0 | 4.991 | 0 ± 0 | 5.096 | 0 ± 0 | 5.004 | 0.035 ± 0 | 1.545 | 1.473 |
|  | ***k*_catC_** | -0.011 ± 0 | 5.083 | -0.011 ± 0 | 5.013 | 0 ± 0 | 4.903 | 0 ± 0 | 4.892 | 0.035 ± 0 | 1.607 | 1.463 |
|  | ***K*_C_** | -0.011 ± 0 | 5.012 | -0.011 ± 0 | 5.038 | 0 ± 0 | 5.065 | 0 ± 0 | 5.043 | 0.035 ± 0 | 1.542 | 1.483 |
|  | ***K*_C_^air^** | -0.011 ± 0 | 5.083 | -0.011 ± 0 | 5.004 | 0 ± 0 | 5.031 | 0 ± 0 | 5.02 | 0.035 ± 0 | 1.644 | 1.567 |
|  | ***K*_O_** | -0.012 ± 0 | 4.919 | -0.012 ± 0 | 4.855 | 0 ± 0 | 5.004 | 0 ± 0 | 4.96 | 0.035 ± 0 | 1.567 | 1.435 |

**Table S3.** The percent of 100,000 arbitrary traits which exhibited a phylogenetic signal that was comparable or stronger than that observed in the real rubisco kinetic data. Arbitrary traits were generated by randomly re-shuffling real rubisco kinetic trait data across species on the phylogenetic tree (table S1) or by randomly simulating data with the same distributional properties as each real rubisco kinetic trait (table S2). Phylogenetic signal is measured using five signal detection methods. Data are shown from the analysis of all angiosperms and C_3_ angiosperms, respectively.

| **Data** | **Arbitrary Trait** | **Re-shuffled** | | | | | **Simulated** | | | | |
| --- | --- | --- | --- | --- | --- | --- | --- | --- | --- | --- | --- |
|  |  | ***Cmean*** | ***I*** | ***K*** | ***K**** | ***Lambda*** | ***Cmean*** | ***I*** | ***K*** | ***K**** | ***Lambda*** |
| **All Angiosperms** | ***S*_C/O_** | 0.001 | 0.001 | 7.087 | 8.279 | 0.010 | 0.001 | 0.001 | 2.980 | 3.613 | 0.001 |
|  | ***k*_catC_** | 0.001 | 0.013 | 0.005 | 0.004 | 0.001 | 0.001 | 0.023 | 0.006 | 0.005 | 0.001 |
|  | ***K*_C_** | 0.001 | 0.005 | 0.204 | 0.108 | 0.001 | 0.003 | 0.009 | 0.224 | 0.103 | 0.001 |
|  | ***K*_C_^air^** | 0.022 | 0.007 | 4.740 | 3.056 | 0.738 | 0.018 | 0.004 | 4.674 | 2.998 | 0.914 |
|  | ***K*_O_** | 24.942 | 85.915 | 96.296 | 95.187 | 26.779 | 25.116 | 85.014 | 98.313 | 97.534 | 25.825 |
| **C_3_ Angiosperms** | ***S*_C/O_** | 0.001 | 0.001 | 40.118 | 38.530 | 0.455 | 0.001 | 0.001 | 29.326 | 26.758 | 0.021 |
|  | ***k*_catC_** | 0.001 | 0.033 | 0.434 | 0.331 | 0.001 | 0.001 | 0.045 | 0.512 | 0.384 | 0.001 |
|  | ***K*_C_** | 0.001 | 0.001 | 3.246 | 2.306 | 0.001 | 0.001 | 0.001 | 3.296 | 2.337 | 0.001 |
|  | ***K*_C_^air^** | 0.001 | 0.001 | 1.423 | 0.965 | 0.001 | 0.001 | 0.001 | 1.304 | 0.869 | 0.001 |
|  | ***K*_O_** | 0.012 | 0.648 | 47.350 | 42.136 | 0.081 | 0.007 | 0.614 | 46.755 | 41.467 | 0.095 |

**Table S4.** The phylogenetic signal strength (mean ± S.E.) and percent of significant tests in arbitrary Brownian motion traits distributed across randomly simulated phylogenetic trees using 100 trait replicates for each of 100,000 tree replicates. Phylogenetic signal is measured using five signal detection methods. Data are shown from the analysis which uses the same total number of species as all angiosperms and C_3_ angiosperms in the rubisco kinetic dataset, respectively.

| **Number of species** | ***Cmean*** | | ***I*** | | ***K*** | | ***K**** | | ***Lambda*** | | **Consensus** |
| --- | --- | --- | --- | --- | --- | --- | --- | --- | --- | --- | --- |
|  | ***Stat*** | ***% Sig*** | ***Stat*** | ***% Sig*** | ***Stat*** | ***% Sig*** | ***Stat*** | ***% Sig*** | ***Stat*** | ***% Sig*** | ***% Sig*** |
| **All Angiosperms** | 0.681 ± 0 | 100.000 | 0.121 ± 0 | 100.000 | 1.000 ± 0 | 99.885 | 1.000 ± 0 | 100.000 | 1.012 ± 0 | 100.000 | 100.000 |
| **C_3_ Angiosperms** | 0.652 ±0 | 99.999 | 0.128 ±0 | 99.997 | 1.000 ±0 | 99.717 | 1.000 ± 0 | 100.000 | 1.009 ±0 | 99.991 | 99.999 |

**Table S5.** The phylogenetic signal strength and associated significance level in rubisco kinetic traits when omitting species in the *Oryza*, *Aegilops* and *Triticum* clades. Phylogenetic signal is measured using five signal detection methods. Data are shown from the analysis of all angiosperms and C_3_ angiosperms, respectively.

NOTE. Statistics are rounded to three decimal places and significance values are represented as α levels, where α = 0.001 if P < 0.001, α = 0.01 if 0.001 < P < 0.01, α = 0.05 if 0.01 < P < 0.05, and α = ns if P > 0.05.

| **Data** | ***Kinetic Trait*** | ***Cmean*** | | ***I*** | | ***K*** | | ***K**** | | ***Lambda*** | |
| --- | --- | --- | --- | --- | --- | --- | --- | --- | --- | --- | --- |
|  |  | ***Stat*** | ***α*** | ***Stat*** | ***α*** | ***Stat*** | ***α*** | ***Stat*** | ***α*** | ***Stat*** | ***α*** |
| **All Angiosperms** | ***S*_C/O_** | 0.527 | 0.001 | 0.477 | 0.001 | 0.002 | 0.05 | 0.002 | 0.05 | 0.881 | 0.001 |
|  | ***k*_catC_** | 0.320 | 0.001 | 0.241 | 0.001 | 0.003 | 0.001 | 0.004 | 0.001 | 0.938 | 0.01 |
|  | ***K*_C_** | 0.285 | 0.001 | 0.284 | 0.001 | 0.003 | 0.01 | 0.004 | 0.01 | 0.659 | 0.001 |
|  | ***K*_C_^air^** | 0.234 | 0.001 | 0.268 | 0.001 | 0.001 | ns | 0.001 | ns | 0.263 | ns |
|  | ***K*_O_** | 0.045 | ns | -0.056 | ns | 0 | ns | 0 | ns | 0 | ns |
| **C_3_ Angiosperms** | ***S*_C/O_** | 0.567 | 0.001 | 0.532 | 0.001 | 0.002 | ns | 0.002 | ns | 0.816 | 0.001 |
|  | ***k*_catC_** | 0.353 | 0.001 | 0.259 | 0.001 | 0.002 | ns | 0.003 | ns | 0.762 | 0.01 |
|  | ***K*_C_** | 0.454 | 0.001 | 0.402 | 0.001 | 0.004 | 0.05 | 0.005 | 0.05 | 0.946 | 0.001 |
|  | ***K*_C_^air^** | 0.401 | 0.001 | 0.370 | 0.001 | 0.003 | ns | 0.003 | ns | 0.940 | 0.001 |
|  | ***K*_O_** | 0.346 | 0.001 | 0.241 | 0.001 | 0.008 | 0.05 | 0.009 | 0.05 | 0.804 | 0.001 |

**Table S6.** The phylogenetic signal strength and associated significance level in rubisco kinetic traits when assessed using phylogenetic trees inferred from the coding sequence of *matk*, *ndhF*, or *atpB* genes. Phylogenetic signal is measured using five signal detection methods. Data are shown from the analysis of all angiosperms and C_3_ angiosperms, respectively.

NOTE. Statistics are rounded to three decimal places and significance values are represented as α levels, where α = 0.001 if P < 0.001, α = 0.01 if 0.001 < P < 0.01, α = 0.05 if 0.01 < P < 0.05, and α = ns if P > 0.05.

| **Data** | ***Kinetic Trait*** | ***Cmean*** | | ***I*** | | ***K*** | | ***K**** | | ***Lambda*** | |
| --- | --- | --- | --- | --- | --- | --- | --- | --- | --- | --- | --- |
|  |  | ***Stat*** | ***α*** | ***Stat*** | ***α*** | ***Stat*** | ***α*** | ***Stat*** | ***α*** | ***Stat*** | ***α*** |
| **All Angiosperms *matK***  **(*n* = 85)** | ***S*_C/O_** | 0.327 | 0.001 | 0.267 | 0.001 | 0.001 | 0.01 | 0.001 | 0.05 | 0.855 | 0.001 |
|  | ***k*_catC_** | 0.335 | 0.001 | 0.21 | 0.01 | 0.001 | 0.01 | 0.001 | 0.001 | 0.845 | 0.05 |
|  | ***K*_C_** | 0.097 | ns | 0.106 | ns | 0 | ns | 0 | ns | 0 | ns |
|  | ***K*_C_^air^** | 0.095 | ns | 0.122 | 0.05 | 0 | ns | 0 | ns | 0 | ns |
|  | ***K*_O_** | -0.058 | ns | -0.106 | ns | 0 | ns | 0 | ns | 0 | ns |
| **All Angiosperms *ndhF***  **(*n* = 76)** | ***S*_C/O_** | 0.389 | 0.001 | 0.313 | 0.001 | 0 | 0.05 | 0 | ns | 0.836 | 0.001 |
|  | ***k*_catC_** | 0.429 | 0.001 | 0.196 | 0.01 | 0.001 | 0.01 | 0.001 | 0.01 | 0.986 | 0.001 |
|  | ***K*_C_** | 0.131 | 0.05 | 0.052 | ns | 0 | ns | 0 | ns | 0.01 | ns |
|  | ***K*_C_^air^** | 0.171 | 0.05 | 0.095 | ns | 0 | ns | 0 | ns | 0.001 | ns |
|  | ***K*_O_** | -0.116 | ns | -0.142 | ns | 0 | ns | 0 | ns | 0 | ns |
| **All Angiosperms *atpB***  **(*n* = 63)** | ***S*_C/O_** | 0.273 | 0.001 | 0.141 | 0.05 | 0 | ns | 0 | ns | 0.866 | 0.001 |
|  | ***k*_catC_** | 0.365 | 0.001 | 0.225 | 0.01 | 0.001 | 0.001 | 0.002 | 0.001 | 0.975 | 0.001 |
|  | ***K*_C_** | 0.274 | 0.001 | 0.205 | 0.01 | 0 | 0.01 | 0.001 | 0.01 | 0.894 | 0.05 |
|  | ***K*_C_^air^** | 0.205 | 0.01 | 0.175 | 0.05 | 0.001 | 0.001 | 0.001 | 0.001 | 0.796 | ns |
|  | ***K*_O_** | 0.130 | 0.05 | 0.009 | ns | 0 | ns | 0 | ns | 0.122 | ns |
| **C_3_ Angiosperms *matK***  **(*n* = 69)** | ***S*_C/O_** | 0.308 | 0.001 | 0.262 | 0.01 | 0.001 | 0.05 | 0.001 | 0.05 | 0.809 | 0.001 |
|  | ***k*_catC_** | 0.396 | 0.001 | 0.283 | 0.001 | 0.001 | 0.01 | 0.001 | 0.01 | 0.884 | 0.001 |
|  | ***K*_C_** | 0.408 | 0.001 | 0.29 | 0.001 | 0 | 0.05 | 0 | 0.05 | 0.865 | 0.001 |
|  | ***K*_C_^air^** | 0.355 | 0.001 | 0.257 | 0.01 | 0 | 0.05 | 0.001 | 0.05 | 0.903 | 0.001 |
|  | ***K*_O_** | 0.186 | 0.05 | 0.044 | ns | 0 | ns | 0 | ns | 0.376 | ns |
| **C_3_ Angiosperms *ndhF***  **(*n* = 51)** | ***S*_C/O_** | 0.111 | ns | 0.173 | 0.05 | 0 | ns | 0 | ns | 0.467 | 0.01 |
|  | ***k*_catC_** | 0.516 | 0.001 | 0.277 | 0.01 | 0.001 | 0.05 | 0.001 | 0.05 | 0.961 | 0.001 |
|  | ***K*_C_** | 0.419 | 0.001 | 0.207 | 0.01 | 0 | ns | 0 | ns | 0.875 | 0.001 |
|  | ***K*_C_^air^** | 0.437 | 0.001 | 0.241 | 0.01 | 0 | ns | 0 | ns | 0.899 | 0.001 |
|  | ***K*_O_** | 0.048 | ns | 0.076 | ns | 0 | ns | 0 | ns | 0 | ns |
| **C_3_ Angiosperms *atpB***  **(*n* = 52)** | ***S*_C/O_** | 0.145 | 0.05 | 0.051 | ns | 0 | ns | 0 | ns | 0.625 | 0.001 |
|  | ***k*_catC_** | 0.51 | 0.001 | 0.327 | 0.001 | 0.001 | 0.001 | 0.001 | 0.001 | 0.966 | 0.001 |
|  | ***K*_C_** | 0.364 | 0.001 | 0.249 | 0.01 | 0 | 0.01 | 0.001 | 0.01 | 0.911 | 0.001 |
|  | ***K*_C_^air^** | 0.34 | 0.001 | 0.253 | 0.01 | 0 | 0.01 | 0.001 | 0.001 | 0.949 | 0.001 |
|  | ***K*_O_** | 0.1 | ns | 0.001 | ns | 0 | ns | 0 | ns | 0.291 | ns |

**Table S7.** The phylogenetic signal strength and associated significance level in rubisco kinetic traits when considering only the subset of rubisco characterized in Orr *et al.*, (2016). Phylogenetic signal is measured using five signal detection methods. Data are shown from the analysis of all angiosperms and C_3_ angiosperms, respectively.

NOTE. Statistics are rounded to three decimal places and significance values are represented as α levels, where α = 0.001 if P < 0.001, α = 0.01 if 0.001 < P < 0.01, α = 0.05 if 0.01 < P < 0.05, and α = ns if P > 0.05. s

| **Data** | ***Kinetic Trait*** | ***Cmean*** | | ***I*** | | ***K*** | | ***K**** | | ***Lambda*** | |
| --- | --- | --- | --- | --- | --- | --- | --- | --- | --- | --- | --- |
|  |  | ***Stat*** | ***α*** | ***Stat*** | ***α*** | ***Stat*** | ***α*** | ***Stat*** | ***α*** | ***Stat*** | ***α*** |
| **All Angiosperms**  **(*n* = 69)** | ***S*_C/O_** | 0.183 | 0.05 | 0.114 | 0.05 | 0.001 | ns | 0.001 | ns | 0.274 | 0.05 |
|  | ***k*_catC_** | 0.345 | 0.001 | 0.241 | 0.001 | 0.002 | ns | 0.003 | ns | 0.906 | 0.001 |
|  | ***K*_C_** | 0.444 | 0.001 | 0.249 | 0.001 | 0.002 | ns | 0.003 | ns | 0.94 | 0.001 |
|  | ***K*_C_^air^** | 0.384 | 0.001 | 0.21 | 0.001 | 0.001 | ns | 0.002 | ns | 0.883 | 0.001 |
|  | ***K*_O_** | 0.247 | 0.01 | 0.111 | 0.05 | 0.009 | 0.05 | 0.01 | 0.05 | 0.543 | 0.05 |
| **C_3_ Angiosperms**  **(*n* = 64)** | ***S*_C/O_** | 0.284 | 0.001 | 0.186 | 0.01 | 0.001 | ns | 0.001 | ns | 0.37 | 0.001 |
|  | ***k*_catC_** | 0.403 | 0.001 | 0.282 | 0.001 | 0.002 | ns | 0.003 | ns | 0.874 | 0.001 |
|  | ***K*_C_** | 0.339 | 0.001 | 0.278 | 0.001 | 0.002 | ns | 0.002 | ns | 0.928 | 0.001 |
|  | ***K*_C_^air^** | 0.293 | 0.001 | 0.223 | 0.001 | 0.001 | ns | 0.001 | ns | 0.819 | 0.01 |
|  | ***K*_O_** | 0.183 | 0.05 | 0.113 | 0.05 | 0.006 | 0.05 | 0.008 | 0.05 | 0.464 | ns |

**Table S8.** The phylogenetic signal strength and associated significance level in rubisco kinetic traits after a reduction of 13 mol mol^−1^ was applied to *S*_C/O_ values in *Limonium* species. Phylogenetic signal is measured using five signal detection methods. Data are shown from the analysis of all angiosperms and C_3_ angiosperms, respectively.

NOTE. Statistics are rounded to three decimal places and significance values are represented as α levels, where α = 0.001 if P < 0.001, α = 0.01 if 0.001 < P < 0.01, α = 0.05 if 0.01 < P < 0.05, and α = ns if P > 0.05.

| **Data** | ***Kinetic Trait*** | ***Cmean*** | | ***I*** | | ***K*** | | ***K**** | | ***Lambda*** | |
| --- | --- | --- | --- | --- | --- | --- | --- | --- | --- | --- | --- |
|  |  | ***Stat*** | ***α*** | ***Stat*** | ***α*** | ***Stat*** | ***α*** | ***Stat*** | ***α*** | ***Stat*** | ***α*** |
| **All Angiosperms** | ***S*_C/O_** | 0.396 | 0.001 | 0.298 | 0.001 | 0.001 | ns | 0.001 | ns | 0.818 | 0.001 |
|  | ***k*_catC_** | 0.350 | 0.001 | 0.224 | 0.001 | 0.003 | 0.001 | 0.003 | 0.001 | 0.968 | 0.001 |
|  | ***K*_C_** | 0.282 | 0.001 | 0.248 | 0.001 | 0.001 | 0.01 | 0.002 | 0.01 | 0.902 | 0.001 |
|  | ***K*_C_^air^** | 0.234 | 0.001 | 0.242 | 0.001 | 0.001 | 0.05 | 0.001 | 0.05 | 0.392 | ns |
|  | ***K*_O_** | 0.032 | ns | -0.07 | ns | 0 | ns | 0 | ns | 0 | ns |
| **C_3_ Angiosperms** | ***S*_C/O_** | 0.363 | 0.001 | 0.255 | 0.001 | 0 | ns | 0 | ns | 0.678 | 0.001 |
|  | ***k*_catC_** | 0.387 | 0.001 | 0.234 | 0.001 | 0.002 | 0.01 | 0.002 | 0.01 | 0.913 | 0.001 |
|  | ***K*_C_** | 0.449 | 0.001 | 0.341 | 0.001 | 0.001 | 0.05 | 0.001 | 0.05 | 0.948 | 0.001 |
|  | ***K*_C_^air^** | 0.398 | 0.001 | 0.317 | 0.001 | 0.001 | 0.05 | 0.002 | 0.01 | 0.947 | 0.001 |
|  | ***K*_O_** | 0.279 | 0.001 | 0.167 | 0.01 | 0 | ns | 0 | ns | 0.743 | 0.001 |

**Table S9.** The phylogenetic signal strength and associated significance level in rubisco kinetic traits when omitting species in *Limonium* clade. Phylogenetic signal is measured using five signal detection methods. Data are shown from the analysis of all angiosperms and C_3_ angiosperms, respectively.

NOTE. Statistics are rounded to three decimal places and significance values are represented as α levels, where α = 0.001 if P < 0.001, α = 0.01 if 0.001 < P < 0.01, α = 0.05 if 0.01 < P < 0.05, and α = ns if P > 0.05.

| **Data** | ***Kinetic Trait*** | ***Cmean*** | | ***I*** | | ***K*** | | ***K**** | | ***Lambda*** | |
| --- | --- | --- | --- | --- | --- | --- | --- | --- | --- | --- | --- |
|  |  | ***Stat*** | ***α*** | ***Stat*** | ***α*** | ***Stat*** | ***α*** | ***Stat*** | ***α*** | ***Stat*** | ***α*** |
| **All Angiosperms** | ***S*_C/O_** | 0.294 | 0.001 | 0.187 | 0.001 | 0 | ns | 0 | ns | 0.753 | 0.001 |
|  | ***k*_catC_** | 0.339 | 0.001 | 0.205 | 0.001 | 0.003 | 0.001 | 0.003 | 0.001 | 0.97 | 0.001 |
|  | ***K*_C_** | 0.185 | 0.01 | 0.164 | 0.01 | 0.001 | 0.01 | 0.001 | 0.01 | 0.825 | ns |
|  | ***K*_C_^air^** | 0.152 | 0.01 | 0.172 | 0.01 | 0.001 | ns | 0.001 | 0.05 | 0 | ns |
|  | ***K*_O_** | -0.002 | ns | -0.101 | ns | 0 | ns | 0 | ns | 0 | ns |
| **C_3_ Angiosperms** | ***S*_C/O_** | 0.220 | 0.01 | 0.089 | ns | 0 | ns | 0 | ns | 0.367 | 0.001 |
|  | ***k*_catC_** | 0.381 | 0.001 | 0.215 | 0.01 | 0.002 | 0.01 | 0.002 | 0.01 | 0.924 | 0.001 |
|  | ***K*_C_** | 0.341 | 0.001 | 0.211 | 0.01 | 0.001 | ns | 0.001 | 0.05 | 0.943 | 0.001 |
|  | ***K*_C_^air^** | 0.299 | 0.001 | 0.203 | 0.01 | 0.001 | 0.05 | 0.002 | 0.05 | 0.944 | 0.001 |
|  | ***K*_O_** | 0.225 | 0.01 | 0.109 | 0.05 | 0 | ns | 0 | ns | 0.726 | 0.01 |
